# Discrete synaptic events induce global oscillations in balanced neural networks

**DOI:** 10.1101/2023.11.12.566734

**Authors:** Denis S. Goldobin, Matteo di Volo, Alessandro Torcini

## Abstract

Neural dynamics is triggered by discrete synaptic inputs of finite amplitude. However, the neural response is usually obtained within the diffusion approximation (DA) representing the synaptic inputs as Gaussian noise. We derive a mean-field formalism encompassing synaptic shot-noise for sparse balanced networks of spiking neurons. For low (high) external drives (synaptic strenghts) irregular global oscillations emerge via continuous and hysteretic transitions, correctly predicted by our approach, but not from the DA. These oscillations display frequencies in biologically relevant bands.

## Introduction

In several contexts the discrete nature of stochastic events should be taken into account to correctly predict the system dynamics. A typical example is represented by shot-noise, which is conveyed by pulses and is therefore discontinous, at variance with white noise, which is associated to thermal fluctuations and is continuous [1]. The inclusion of shot-noise is fundamental to fully characterize the emergent phenomena in many fields of physics ranging from mesoscopic conductors [2] to driven granular gases [3].

The discrete nature of the events is an innate characteristic also of the neural dynamics, where a neuron receives inputs from other neurons via electrical pulses, termed post-synaptic potentials (PSPs). The PSPs stimulating a neuron in the cortex are usually assumed to be uncorrelated with small amplitudes and high arrival rates. Therefore the synaptic inputs can be treated as a continuous Gaussian process and the neural dynamics can be examined at a mean-field level within the framework of the Diffusion Approximation (DA) [4, 5]. In this context, the theory of dynamical balance of excitation and inhibition [6–8] represents one of the most successfull results able to explain some of the main aspects of cortical dynamics [9].

However, several experiments have shown that rare PSPs of large amplitude can have a fundamental impact on the cortical activity [10, 11] and that synaptic weight distributions display a long tail towards large amplitudes [12–14].

Furthermore, networks of inhibitory neurons with low connectivity (in-degree *K*≃ 30 − 80) have been identified in the cat visual cortex [15] and in the rat hippocampus [16] and the latter are believed to be at the origin of collective oscillations (COs) in the *γ*-band [17]. Recent experiments have also shown that the cortical connections are definitely more sparse in primate when compared to mouse [18].

These experimental evidences call for the development of a mean-field formalism able to incorporate the effect of discrete synaptic events for diluted random networks. Population based formalisms taking into account the discrete nature of the synaptic events have been previously developed for Integrate-and-Fire models [19–22]. However, such approaches are limited to stationary solutions and they cannot describe the emergence of oscillatory behaviours.

In this Letter, we introduce a *complete* mean-field (CMF) approach for balanced neural networks [6], taking into account the sparsness of the network and the discreteness of the synaptic pulses, able to reproduce all the possible dynamical states. For simplicity, but without any loss of generality, we consider inhibitory balanced networks subject to an external excitatory drive [23–26].

Firstly, we illustrate that the DA cannot capture oscillatory behaviours emerging for sufficiently low in-degree in spiking neural networks by considering conductance- and current-based neuronal models. However, this regime is correctly reproduced by a mean-field approach whenever the sparse and discrete synaptic inputs are taken in account. Furthermore, for Quadratic Integrate-and-Fire (QIF) [27, 28] neuronal network via the CMF approach we obtain a complete bifurcation diagram encompassing asynchronous and oscillatory regimes. In particular, for sufficiently low (large) excitatory drive (synaptic amplitudes) the CMF reveals bifurcations from the asynchronous irregular (AI) to the oscillatory irregular (OI) regime as well as a region of coexistence of these two phases not captured by the DA [29]. Exact event-driven simulations of large QIF networks confirm the sub- and super-critical Hopf bifurcations predicted within the CMF theory. Furthermore, for low in-degrees COs in biologically relevant frequency bands (from *δ* to *γ* band) are observable [30].

### The balanced network

As a prototype of a dynamically balanced system we consider a sparse inhibitory network made of *N* pulse-coupled neurons whose membrane potential evolves according to the equations

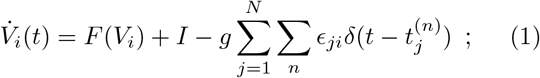

where *I* is an external DC current, *g* the synaptic coupling, and the last term represents the inhibitory synaptic current. The latter is the linear superposition of instantaneous inhibitory PSPs emitted at times 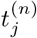 from the pre-synaptic neurons connected to neuron *i. ϵ*_*ji*_ is the adjacency matrix of the random network with entries 1 (0) if the the connection from node *j* to *i* exists (or not), and we assume the same in-degree *K* = _*j*_ *ϵ*_*ji*_ for all neurons. We consider two paradigmatic models of spiking neuron: the quadratic integrate-and-fire (QIF) with *F* (*V*) = *V* ^2^ [25–27, 31, 32], which is a current-based model of class I excitability; and the Morris-Lecar (ML) [33], a conductance-based model representing a class II excitable membrane [34]. The DC current and th <u>e sy</u>naptic coupare assumed to scale as 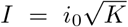 and 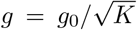 as usually done in order to ensure a self-sustained balanced state for sufficiently large in-degrees [6, 7, 24–26, 35]. The times (frequencies) are reported in physical units by assuming a membrane time constant *τ*_*m*_ = 10 ms.

### Mean-field description

For a sufficiently sparse network, the spike trains emitted by *K* pre-synaptic neurons can be assumed to be uncorrelated and Poissonian [8, 23], therefore the mean-field dynamics of a generic neuron can be represented in terms of following Langevin equation:

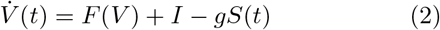

where *S*(*t*) is a Poissonian train of *δ*-spikes with rate *R*(*t*) = *Kν*(*t*), and *ν*(*t*) is the population firing rate self-consistently estimated. Usually the Poissonian spike trains are appro<u>xim</u>ated within the the DA [5, 36] as *S*(*t*) = *R*(*t*) + *R*(*t*)*ξ*(*t*), where *ξ*(*t*) is a Gaussian white noise term. However, this approximation can fail to reproduce fundamental aspects of the neural dynamics. Indeed, as shown in Fig. 1(a) for a sparse ML network, by employing the DA in (2) one obtains an asynchronous dynamics (blue curve), while the correct network evolution, characterized by global oscillations with frequency *f*_*C*_ ≃ 18 Hz (black dots), can be recovered only by explicitely taking into account the Poissonian spike trains in (2) (red line).

**FIG. 1.**
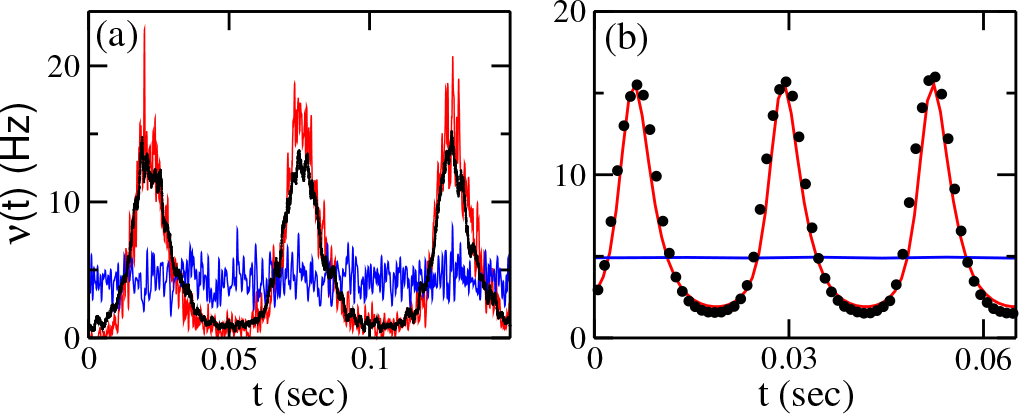
Population firing rate *ν*(*t*) versus time for ML (a) and QIF (b) models: blue (red) lines refer to diffusive (shot-noise) MF results and black circles to network simulations. For the ML the MF shot-noise and DA results have been obtained by integrating the Langevin equation (2), while for the QIF by integrating (3) and (4), respectively: more details in [34]. The parameters for the ML model are *K* = 20, *i*_0_ = 0.1, *g*_0_ = 5 and network size *N* = 20000, the other parameters are reported in the supplemental material [34]. For the QIF model *K* = 200, *i*_0_ = 0.16, *g*_0_ = 4 and *N* = 80000.

In the mean-field framework the population dynamics is usually described in terms of the membrane potential probability distribution function (PDF) *P* (*V, t*), whose time evolution is given for the QIF model by the following continuity equation

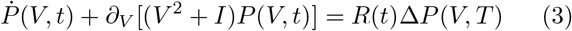

with boundary condition lim_*V →∞*_ *V* ^2^*P* (*V, t*) = *ν*(*t*) and where Δ*P* (*V, T*) = [*P* (*V* ^+^, *t*) − *P* (*V, t*)] with *V* ^+^ = *V* +*g*. By assuming that *g* is sufficiently small we can expand the latter term as 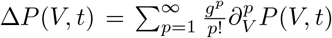; and by limiting to the first two terms in this expansion we recover the DA corresponding to the following Fokker-Planck Equation (FPE) [37]

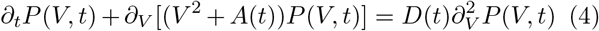

where 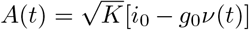 and 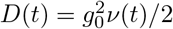. The DA can give uncorrect predictions for the QIF model, as well. Indeed as shown in Fig. 1 (b) the network dynamics is oscillatory with *f*_*c*_ ≃ 40 Hz (black circles) : an evolution correctly captured by the MF equation (3) (red line), while the FPE (4) converges to a a stable fixed point (blue curve), Therefore to reproduce the collective dynamical regimes observable in the network it is necessary to consider the complete continuity equation (3). In this respect we have developed a CMF formalism encompassing synaptic shot-noise to identify the various possible regimes displayed by (3) and to analyse their stability.

The QIF model evolution can be transformed in that of a phase oscillator, the so-called *θ*-neuron [27, 38], by introducing the phase variable *θ* = 2 arctan *V*. However, this transformation has the drawback that even uncoupled neurons are associated to a non flat PDF of the phases, thus rendering quite difficult or even unfeasible to identify asynchronous regimes with respect to partially synchronized ones in noisy enviroments [39, 40]. A more appropriate phase transformation to analyse the synchron<u>iz</u>ation phenomena is the following *ψ* = 2 arctan 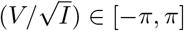, which leads to a uniformly rotating phase in the absence of incoming pulses for suprathreshold neurons with *I >* 0 [34].

By con<u>s</u>idering the phase PDF 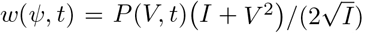 Eq. (3) can be rewritten in terms of t the so-called Kuramoto–Daido order parameters *z*_*n*_ [41, 42] by expanding in Fourier space the PDF as 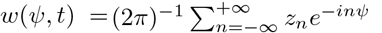 with *z*_0_ = 1 and 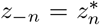. After laborious but straightforward calculations, one obtains the following evolution equations

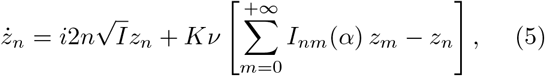

where *n* = 1, 2, 3, …, 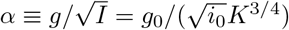,

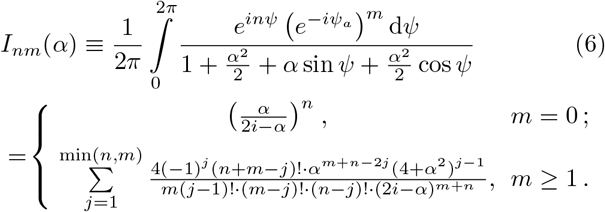

The firing rate can be self-consistently determined by the flux at the firing threshold 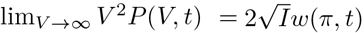, as follows

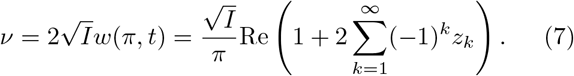

The dynamics of the system (5,7) is controlled by only two parameters: *K* and *α*. Thus, we can limit to derive a bidimensional phase diagram in the parameter plane 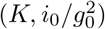, that will comprehensively cover the entire diversity of the macroscopic regimes observable in the network. In particular, we have estimated the stationary solutions of Eqs. (5,7) by truncating the Fourier expansion in (5) to *M* ≥ 100 modes in order to guarantee a numerical accuracy of *𝒪* (10^*−*12^) for all the parameter values. The linear stability of the asynchronous state has allowed us to identify a HB line where the oscillatory dynamics emerges: this is reported as a orange line in Fig. 2 (a) together with the super-critical HB line obtained within the DA (black solid line) previously reported in [43]. At variance with the DA the HBs induced by the shot-noise can be either super(solid orange line) or subcritical (dashed orange line), thus allowing for regions where asynchronous and oscillatory regimes can coexist, see Fig. 2 (b). Furthermore while for the DA the oscillatory dynamics is observable only for sufficiently large in-degree *K* ≥ *K*_*min*_ ≃ 220, by taking into account the discrete nature of the synaptic events COs may emerge even for extremely small in-degrees. Furthermore, the asynchronous regime is always unstable for sufficiently small *i*_0_ (large *g*_0_) : namely, for for 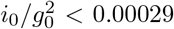. A peculiarity of the shot-noise results is that the HB line is re-entrant, thus in a certain range of 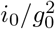we can have asynchronous dynamics only in a finite interval of in-degrees (as shown in Fig. 2 (c)).

**FIG. 2.**
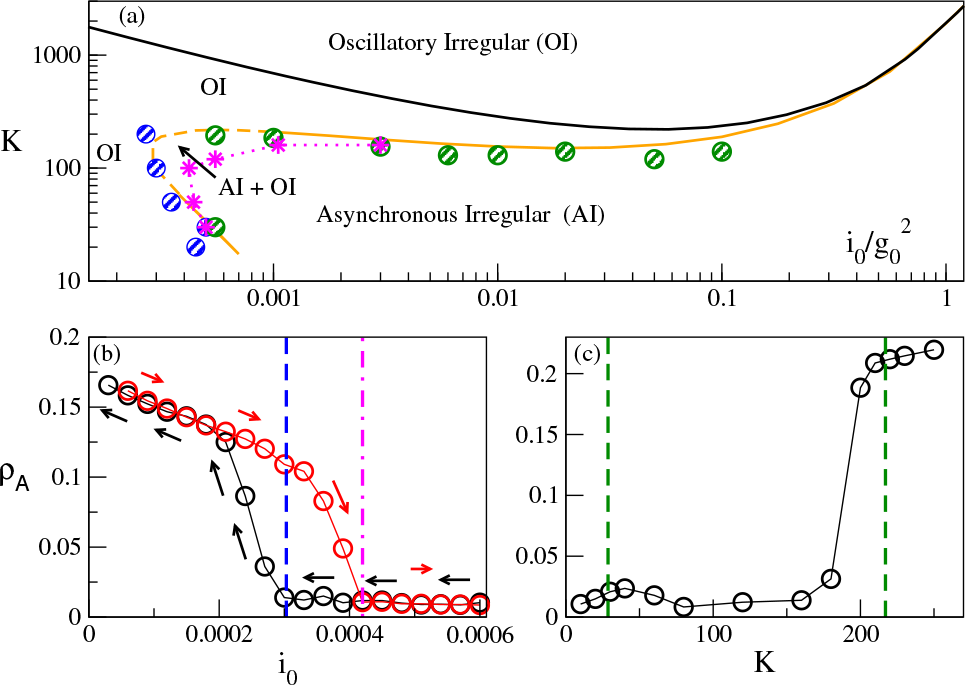
(a) Phase diagram for the QIF network in the plane 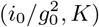: the black solid line is the super-critical HB line obtained within the DA; the orange solid (dashed) line is the super- (sub-) critical HB line given by the CMF; the symbols refer to numerical estimations of the HBs and Saddle-Node Bifurcations (SNBs). The green (blue) circles denote HBs obtained by performing quasi-adiabatic simulations by varying *K* (*i*_0_) for constant *i*_0_ (*K*) values; the magenta stars indicate SNBs. For more details see [34]. (b-c) Average order parameter *ρ*_*A*_ versus *i*_0_ (*K*) for quasi-adiabatic simulations : black circles refer to decreasing (increasing) *i*_0_ (*K*), while red ones to increasing (decreasing) *i*_0_ (*K*). The blue dashed line in (b) denotes the sub-critical HB given by the CMF and the magenta dot-dashed line to numerically estimated SNB; the two green dashed lines in (c) indicate the HBs given by the CMF. The values of *ρ*_*A*_ in panel (b) (panel (c)) refer to *K* = 100 (*i*_0_ = 0.00055) averaged over 5 network realizations, with *N* = 80000, for a time interval *t* = 30 following a transient of 20 s. All data refer to *g*_0_ = 1.

### Network Simulations

In order to verify the CMF predictions we have performed essentially exact numerical simulations of the QIF network by employing a fast event-driven integration scheme [44], which allowed us to follow the network dynamics for long times, up to 50−100 sec, for system of sizes *N* = 10000 − 80000 [34]. In particular, to characterize the macroscopic evolution of the network we measured the following indicator [45]

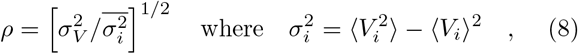

and *σ*_*V*_ is the standard deviation of the mean membranepotential 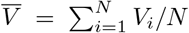, with 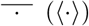 denoting an ensemble (a time) average. A coherent macroscopic activity is associated with a finite value of *ρ* (perfect synchrony corresponds to *ρ* ≡ 1), while an asy<u>nc</u>hronous dynamics to a vanishingly small 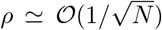. A finite size analysis of the order parameter *ρ*_*A*_ averaged over several different network realizations has allowed us to identify the HBs and the Saddle-Node Bifurcations (SNBs) of limit cycles displayed in Fig. 2. In particular, in Fig. 2 (a) green (blue) circles refer to HBs identified via quasi-adiabatic simulations by varying *K* (*i*_0_) for constant *i*_0_ (*K*) values; while the magenta stars indicate SNBs. Numerical simulations are in good agreement with the CMF results and allowed us also the identification of a coexistence region for asynchronous irregular and oscillatory irregular dynamics. By irregular we mean that the microscopic evolution is characterized by fluctuations in the instantaneous firing rates associated to coefficient of variations [46] of *𝒪* (1), as we have verified [23]. A hysteretic transition from AI to OI obtained by varying quasi-adiabatically *i*_0_ is displayed in Fig. 2 (b), the coexistence region can be clearly identified between the sub-critical HB (blue dashed line) and the SNB (magenta dashed line). Furthermore, as shown in Fig. 2 (c) for sufficiently small currents AI states are observables only for intermediate values of the in-degrees (*K* ∈ [50 : 180] in the considered case), bounded by regions at small (*K* ≤ 40) and large (*K* ≥ 200) in-degrees where OI are instead present. The finite-size scaling analysis of *ρ*_*A*_ for this specific case, revealing the different regimes, is reported in Fig. S1 in [34].

At the HBs, COs emerge with a frequency *f* ^*H*^ that is reported as a function of 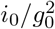 in Fig. 3 (a). The comparison between the results of the CMF approach (solid line) and of network simulations with *N* = 20000 (blue stars) is very good along the whole bifurcation line predicted by the CMF. Furthermore, *f* ^*H*^ covers a wide range of frequencies ranging from 1.77 Hz (*δ* band) to ≃ 100 Hz (*γ* band).

**FIG. 3.**
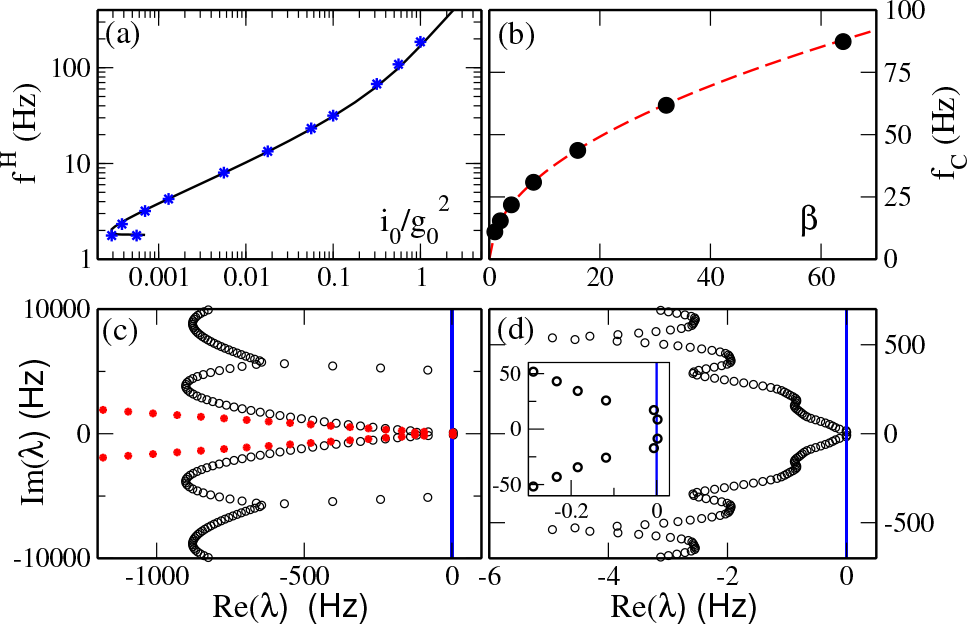
(a) Frequency *f* ^*H*^ of the COs at the HB versus 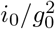: symbols are simulations for *N* = 20000 and the solid line are the CMF results.(b) Frequency *f*_*C*_ of the COs a<u>s</u> a function of the parameter *β*, where *i*_0_ = *β ×* 0.01, *g*_0_ = 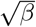, *K* = 200. Circles are network simulation data with *N* = 20<u>0</u>00 and the red dashed line represents the curve *ν*_*CO*_ = 11 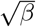 Hz. (c-d) Spectrum of the eigenvalues *{λ*_*i*_ *}* for a stationary solution of system (5,7) for 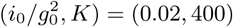 (c), and (0.00055, 10) (d). An enlargement is reported in the inset in (d). Black circles (red stars) refer to the CMF (DA).

As expected by the CMF analysis, the same dyanmics should be observable at fixed *K* by maintaing the ratio 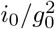 constant. Indeed this is the case, as we have verified by considering a state in the OI regime corresponding to 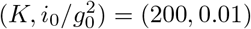 and by varying, as a function of a control paramer *β*, the synaptic coupling and the current as 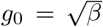 and *i*_0_ = *β* × 0.01, while *K* stays fconstant. We obsvered irregular dynamics characterized by an average 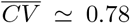 [46] and COs in the whole examined range *β* ∈ [1, 64]. As expected, the only observable variation is in the time scale, that decreases as 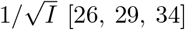, consequently the frequency *f*_*C*_ of the COs grows proportionally to 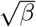, thus one can observe OI dynamics induced by finite amplitude PSP in a wide frequency range by simply varying the parameter *β* (see Fig. 3 (a)).

### Stability of the Asynchronous Regime

#### DA versus CMF

The linearization of the system (5,7) allows us to perform a linear stability analysis of the asynchronous regime, corresponding to a constant firing rate. In particular, we have estimated the corresponding complex spectrum {*λ*_*i*_} : the fixed point is stable whenever *Re λ*_*i*_ *<* 0 ∀*i*. Here we would like to compare the spectra obtained within the DA and the CMF to better understand the origin of the instabilities leading to oscillatory dynamics in presence of microscopic shot-noise. As a first remark, we observe that the DA spectra are characterized besides the most unstable modes, which can give rise to the oscillatory instability, by modes that are strongly damped as shown in Fig. 3 (c). The case shown in Fig. 3(c) refers to a situation where the dynamics is well reproduced within the DA, in this case the DA eigenvalues corresponding to small Im *λ*_*i*_ in proximity of the Hopf instability approximate quite well the CMF spectrum. However, while the CMF eigenvalues appear to saturate at some finite Re *λ* value, the DA ones do not. Despite this difference in this case the collective dynamics of the system is essentially controlled by the two most unstable modes, that pratically coincide within the DA and CMF approaches.

In Fig. 3(d) we report the CMF spectrum for a situation where the OI regime is definitely due to the finitess of the synaptic stimulations and not captured at all by the DA. In this case, we observe that a large part of the eigenmodes are now practically not damped, compare the scales over which Re *λ*_*i*_ varies in Fig. 3(c) and (d). Therefore, we expect that the collective dynamics is no more dominated by only the 2 most unstable modes as usually observable in the DA, but that also the marginally stable or slightly unstable modes will have a role in the coherent dynamics, see the inset of panel (d). In summary, the shot-noise promotes the emergence of weakly damped eigenmodes that have a relevant role in the instability of the asynchronous regime at sufficiently small in-degrees and that are neglected in the DA.

### Conclusions

We have shown that the macroscopic phase-diagram of balanced networks is strongly influenced by the discreteness and the finite amplitude of PSPs. In particular, we have developed a CMF formalism by including Poissonian shot-noise which reproduces quite well the network simulations, at variance with the DA. Our mean-field analysis of the balanced state complements the previous ones, that has been performed in the conxtext of the DA [23] or in the limit *N >> K >>* 1 [6], and adresses some aspects of the neural dynamics not taken into account by the previous analysis. A counterintuitive aspect is the fact that COs can be observed even in extremely sparse inhibitory networks with frequencies in a wide range from 1-2 Hz (*δ*-band) to 100 Hz (*γ*-band). Thus somehow supporting the supposition reported in [17] that *γ*-oscillations in the hippocampus are generated by sub-networks of interneurons with low in-degrees *K* ≃ 30 − 80 [16].

Our analysis has been limited to homogeneous networks, the inclusion of heterogeneity in the mean-field formulation could be probably worked out by assuming Lorentzian distributed heterogeneities which can be analytically integrated [47–49], somehow similarly to what done within the DA in [29].

Quite recently, the effect of shot-noise induced by finite size fluctuations have been analyzed for the macrosocpic evolution of globally coupled populations of QIF neurons [50, 51]. It will be interesting to combine such approach with our to fully understand the relevance of finite-size fluctuations for the dynamics of random sparse networks.

We acknowledge stimulating discussions with Alberto Bacci, Alberto Ferrara, Nina La Miciotta, Lyudmila Klimenko, Gianluigi Mongillo, Simona Olmi, Antonio Politi. D.S.G. acknowledges the support of the CNR Short Term Mobility Programme 2021 for a visit to Istituto dei Sistemi Complessi, Sesto Fiorentino, Italy where part of this work was developed. A.T. received financial support by the Labex MME-DII (Grant No. ANR-11-LBX-0023-01), by CY Generations (Grant No ANR-21-EXES-0008), and together with M.V. by the ANR Project ERMUNDY (Grant No. ANR-18-CE37-0014) and M.V. by the Labex CORTEX (Grant No. ANR-11-LABX-0042) of Université Claude Bernard Lyon 1 and by the the ANR via the Junior Professor Chair in Computational Neurosciences Lyon 1.

## Supporting information

See for for details on the employed neural models, on the integration of the neural networks as well as of the population models, an of the complete m

## References

[1] W. Schottky, Annalen der Physik 362, 541 (1918).

[2] Y. M. Blanter and M. Büttiker, Physics reports 336, 1 (2000).

[3] D. Lucente, M. Viale, A. Gnoli, A. Puglisi, and A. Vulpiani, Physical Review Letters 131, 078201 (2023).

[4] R. Capocelli and L. Ricciardi, Kybernetik 8, 214 (1971).

[5] H. C. Tuckwell, Introduction to theoretical neurobiology: nonlinear and stochastic theories, Vol. 2 (Cambridge University Press, 1988).

[6] C. van Vreeswijk and H. Sompolinsky, Science 274, 1724 (1996).

[7] A. Renart, J. de la Rocha, P. Bartho, L. Hollender, N. Parga, A. Reyes, and K. D. Harris, Science 327, 587 (2010).

[8] N. Brunel, Journal of Computational Neuroscience 8, 183 (2000).

[9] J. Barral and A. D. Reyes, Nature neuroscience 19, 1690 (2016).

[10] S. Song, P. J. Sjöström, M. Reigl, S. Nelson, and D. B. Chklovskii, PLoS biology 3, e68 (2005).

[11] S. Lefort, C. Tomm, J.-C. F. Sarria, and C. C. Petersen, Neuron 61, 301 (2009).

[12] R. Miles, The Journal of Physiology 431, 659 (1990).

[13] B. Barbour, N. Brunel, V. Hakim, and J.-P. Nadal, TRENDS in Neurosciences 30, 622 (2007).

[14] G. Buzsáki and K. Mizuseki, Nature Reviews Neuroscience 15, 264 (2014).

[15] Z. F. Kisvárday, C. Beaulieu, and U. T. Eysel, Journal of comparative neurology 327, 398 (1993).

[16] A. Sik, M. Penttonen, A. Ylinen, and G. Buzsaki, Journal of Neuroscience 15, 6651 (1995).

[17] G. Buzsáki and X.-J. Wang, Annual review of neuroscience 35, 203 (2012).

[18] G. A. Wildenberg, M. R. Rosen, J. Lundell, D. Paukner, D. J. Freedman, and N. Kasthuri, Cell Reports 36 (2021).

[19] M. J. Richardson and R. Swarbrick, Physical review letters 105, 178102 (2010).

[20] R. Iyer, V. Menon, M. Buice, C. Koch, and S. Mihalas, PLoS computational biology 9, e1003248 (2013).

[21] S. Olmi, D. Angulo-Garcia, A. Imparato, and A. Torcini, Scientific reports 7, 1577 (2017).

[22] F. Droste and B. Lindner, Journal of computational neuroscience 43, 81 (2017).

[23] N. Brunel and V. Hakim, Neural computation 11, 1621 (1999).

[24] J. Kadmon and H. Sompolinsky, Phys. Rev. X 5, 041030 (2015).

[25] M. Monteforte and F. Wolf, Phys. Rev. Lett. 105, 268104 (2010).

[26] M. di Volo and A. Torcini, Phys. Rev. Lett. 121, 128301 (2018).

[27] G. B. Ermentrout and N. Kopell, SIAM Journal on Applied Mathematics 46, 233 (1986).

[28] B. Gutkin, in Encyclopedia of computational neuroscience (Springer, 2022) pp. 3412–3419.

[29] M. Di Volo, M. Segneri, D. S. Goldobin, A. Politi, and A. Torcini, Chaos: An Interdisciplinary Journal of Nonlinear Science 32, 023120 (2022).

[30] G. Buzsaki, Rhythms of the Brain (Oxford University Press, 2006).

[31] C. R. Laing, The Journal of Mathematical Neuroscience 8, 1 (2018).

[32] I. Ratas and K. Pyragas, Physical Review E 100, 052211 (2019).

[33] C. Morris and H. Lecar, Biophysical journal 35, 193 (1981).

[34] See Supplemental Material at [URL will be inserted by publisher] for details on the employed neural models, on the integration of the neural networks as well as of the population models, an of the complete mean-field analysis.

[35] A. Litwin-Kumar and B. Doiron, Nat Neurosci 15, 1498 (2012).

[36] R. Capocelli and L. Ricciardi, Kybernetik 8, 214 (1971).

[37] E. Haskell, D. Q. Nykamp, and D. Tranchina, Network: Computation in Neural Systems 12, 141 (2001).

[38] B. Ermentrout, Scholarpedia 3, 1398 (2008), revision #122134.

[39] B. Kralemann, L. Cimponeriu, M. Rosenblum Pikovsky, and R. Mrowka, Physical Review E 76, 055201 (2007).

[40] A. V. Dolmatova, D. S. Goldobin, and A. Pikovsky, Physical Review E 96, 062204 (2017).

[41] Y. Kuramoto, Chemical oscillations, waves, and turbulence, Vol. 19 (Springer Science & Business Media, 2012).

[42] H. Daido, Progress of theoretical physics 88, 1213 (1992).

[43] M. Di Volo, M. Segneri, D. S. Goldobin, A. Politi, and A. Torcini, Chaos: An Interdisciplinary Journal of Non-linear Science 32 (2022).

[44] A. Tonnelier, H. Belmabrouk, and D. Martinez, Neural Computation 19, 3226 (2007).

[45] D. Golomb, Scholarpedia 2, 1347 (2007).

[46] The coefficient of variation cv(i) for the neuron i is the ratio between the standard deviation and the mean of the interspike intervals associated with its firing activity. *CV* is the ensemble average of the single neurons cv(i).

[47] E. Yakubovich, SOVIET PHYSICS JETP 8.

[48] E. Ott and T. M. Antonsen, Chaos: An interdisciplinary journal of nonlinear science 19 (2009).

[49] E. Montbrió, D. Pazó, and A. Roxin, Physical Review X 5, 021028 (2015).

[50] V. V. Klinshov and S. Y. Kirillov, Physical Review E 106, L062302 (2022).

[51] V. Klinshov, P. Smelov, and S. Y. Kirillov, Chaos: An Interdisciplinary Journal of Nonlinear Science 33 (2023).

