## Supplementary material for "Discrete synaptic events induce global oscillations in balanced neural networks": See for for details on the employed neural models, on the integration of the neural networks as well as of the population models, an of the complete m

### Supplemental material : Discrete spiking events induce global oscillations in balanced neural networks

[1] *Institute of Continuous Media Mechanics, Ural Branch of RAS, Acad. Korolev street 1, 614013 Perm, Russia*
[2] *Department of Theoretical Physics, Perm State University, Bukirev street 15, 614990 Perm, Russia*
[3] *Université Claude Bernard Lyon 1, Institut National de la Santé et de la Recherche Médicale, Stem Cell and Brain Research Institute U1208, Bron, France*
[4] *Laboratoire de Physique Théorique et Modélisation, Université de Cergy-Pontoise, CNRS, UMR 8089, 95302 Cergy-Pontoise cedex, France*
[5] *CNR - Consiglio Nazionale delle Ricerche - Istituto dei Sistemi Complessi, via Madonna del Piano 10, I-50019 Sesto Fiorentino, Italy*

(Dated: November 10, 2023)

### S1. THE MODELS

#### A. The Morris-Lecar Model

For the Morris-Lecar model the nonlinear function $F(V)$ entering in eq. (1) in the Letter takes the form

$$F(V) = g_L(E_L - V) + g_{Ca}M_\infty(V)(E_{Ca} - V) + g_K N(V)(E_K - V) + I_c \tag{S1}$$

where the dynamics of $N(v)$ is ruled by the first order ODE

$$\dot{N}(V) = \frac{N_\infty(V) - N(V)}{\tau_N(V)} \quad , \tag{S2}$$

and

$$M_\infty(V) = \frac{1}{2}\Big(1 + \tanh\Big((V - E_1)/E_2\Big)\Big) \quad ; \tag{S3}$$

$$N_\infty(V) = \frac{1}{2}\Big(1 + \tanh\Big((V - E_3)/E_4\Big)\Big) \quad ; \tag{S4}$$

$$\tau_N(V) = 15 \ \mathrm{sech}((V - E_3)/(2E_4)) \quad . \tag{S5}$$

The parameters are chosen in order to have a dynamics corresponding to a Class II excitable membrane as follows: $E_1 = -1.2$, $E_2 = 18$, $E_3 = 2$, $E_4 = 17.4$, $E_L = -60$, $E_K = -80$, $E_{Ca} = 120$, $g_{Ca} = 4.4$, $g_L = 2$, $g_K = 8$, $I_c = 46.15$.

#### B. The Quadratic Integrate-and-Fire Model

As already mentioned in the Letter, the QIF model corresponds to $F(V) = V^2$, where the membrane potential $V \in ]-\infty : +\infty[$. In particular, whenever the membrane potential $V$ reaches infinity, a $\delta$-spike is emitted and instantaneously delivered to the post-synaptic neurons and $V$ is reset to $-\infty$. In absence of synaptic coupling, the QIF model displays excitable (oscillatory) dynamics for $I < 0$ ($I > 0$). Since we consider a purely inhibitory network in order to have non trivial collective dynamics the neurons should be supra-threshold, i.e. with $I > 0$.

### S2. INTEGRATION METHODS

#### A. ML neuronal population

The numerical integration of the ML network model (1) has been done by employing a standard Euler scheme with integration time step $dt = 5 \times 10^{-4}\tau_m$.

For what concerns the integration of the MF Langevin equation (2), we considered $N = 10000$ replica of such equation, corresponding to $N$ *uncoupled* ML neurons. In particular, we integrated such $N$ uncoupled evolution equations (2) by employing an Euler integration scheme with $dt = 5 \times 10^{-4}\tau_m$. The population firing rate $\nu(t)$ entering in eq. (2) at time $t$ is estimated self-consistently by counting the number of spikes $N_{sp}(\Delta t)$ emitted by the $N$ neurons in the preceding time interval of duration $\Delta t = 0.005\tau_m$, as follows

$$\nu(t) = \frac{N_{sp}(\Delta t)}{\Delta t N} \quad . \tag{S6}$$

For the shot-noise case, at the end of every time interval $\Delta t$, every neuron receives a IPSP of finite amplitude $g$ with a probability $K * \nu(t) * \Delta t$ in order to mimick a Poissonian process with rate $K\nu(t)$.

To simulate the Langevin evolution (2) within the DA, we considered $N$ uncoupled neurons whose membrane potential follows the stochastic differential equation

$$\dot{V}(t) = F(V) + I - g_0\sqrt{K}\nu(t) + g_0\sqrt{\nu(t)}\xi(t) \tag{S7}$$

where $\xi$ is Gaussian white noise with zero mean and unitary variance. The firing rate is estimated by the activity of the $N$ neurons via (S6) by performing an average on a sliding time window of duration $\Delta t = 0.005\tau_m$. The integration is performed via an Euler scheme with time step $dt = 5 \times 10^{-4}\tau_m$. An intial transient time $t_t = 10$ s is discarded in all the simulations.

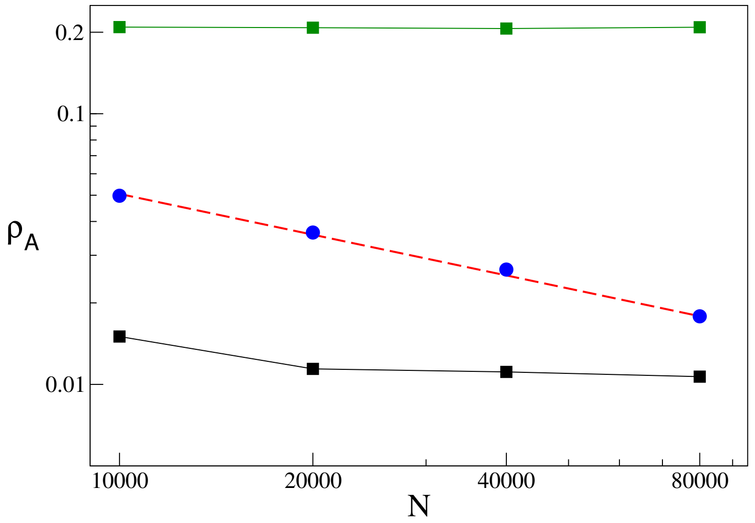


FIG. S1. Average order parameter $\rho_A$ versus the system size $N$ for $K = 10$ (black squares), $K = 60$ (blue circles) and $K = 210$ (grren squares). The solid lines are a guide for the eyes, while the red dashed line denotes a power-law decay $\propto N^{-1/2}$. The values of $\rho_A$ have been averaged over 5-20 network realizations, with $N = 10000 - 80000$, for a time interval $t = 20 - 30$ s following a transient of $t_t = 20 - 30$ s. All data refer to QIF spiking neuronal networks with $i_0 = 0.00055$ and $g_0 = 1$.

##### B. QIF neuronal population

The simulation of the QIF network model (1) has been performed thanks to an exact event-driven method [1], this allows to reach very large system sizes from $N = 10000$ up to $N = 80000$ with $10 \leq K \leq 1000$ for the data in Fig. 1 and Fig. 2.

For what concerns the data reported in Fig 1 (b) in the Letter for the QIF model, the diffusive (shot-noise) MF continuity equation (3) ( FPE (5)) has been integrated with pseudo-spectral time-split methods similar to the ones employed in [2]. In particular, for the Eq. (3) (Eq. (5)) we have transformed the membrane potential to a phase variable defined between $[-\pi : \pi]$ and discretized the phase interval on a grid composed of 128 (512) points, moreover we have employed an integration time step $\Delta t = 1 \times 10^{-4}\tau_m$ ($\Delta t = 1 \times 10^{-5}\tau_m$).

The HBs and SNBs reported in Fig. 2(a) have been obtained by performing finite size analysis of the order parameter $\rho$ obtained during quasi-adiabatic simulations by varying $K$ ($i_0$) for constant $i_0$ ($K$). In particular, the average value of the order parameter $\rho_A$ for each system size $N$ have been obtained by averaging over a time $t = 20 - 30$ s after discarding a transient of $t_t = 20 - 30$ s and over 5 to 20 different network realizations. Analogously to what done in [3], we have classified the dynamical regimes by estimating $\rho_A$ for different network sizes: namely, $N = 10000$, 20000, 40000 and 80000. If the indicator $\rho_A$ decreases as $1/\sqrt{N}$ (saturates to some constant value) then the dynamics is identified as asynchronous (oscillatory) as shown in Fig. S1 for $K = 60$ ($K = 10$ and $K = 210$) at $i_0 = 0.00055$. .

#### S3. MACROSCOPIC DYNAMICS OF LARGE POPULATIONS OF QIF NEURONS

##### A. Derivation of the macroscopic evolution equations

As already mentioned in the Letter, the dynamical evolution of the QIF model can be transformed in that of a phase oscillator, the $\theta$-neuron [4, 5], by introducing the phase variable $\theta = 2 \arctan V$. However, this transformation renders quite difficult to distinguish asynchronous from partially synchronized states. A more suitable phase transformation, able to take into account correctly the phase synchronization phenomena, is the following

$$\psi = 2 \arctan \frac{V}{\sqrt{I}} \in [-\pi, \pi] \ , \qquad V = \sqrt{I} \tan \frac{\psi}{2} \in ] - \infty, +\infty[ \ ,$$

where $I = i_0\sqrt{K}$. Indeed, for $I > 0$ and in the absence of incoming pulses the phase $\psi$ is uniformly rotating with angular velocity $2\sqrt{I}$ at variance with the $\theta$-neuron where even for uncoupled oscillators the velocity depends on the phase.

The PDF $w(\psi)$ for the phase variables $\psi$ is related to the one of the membrane potentials $P(V,t)$ via the relation

$$w(\psi,t) = P(V,t)\frac{I+V^2}{2\sqrt{I}}$$

due to the probability conservation under (nonlinear) transformations of stochastic variables. Therefore, the continuity equation Eq. (3) for $P(V,t)$ :

$$\frac{\partial P(V,t)}{\partial t} = -\frac{\partial}{\partial V}\Big[\big(I+V^2\big)P(V,t)\Big] + K\nu(t)\Big[P(V+g,t) - P(V,t)\Big] , \tag{S8}$$

can be recast as follows for $w(\psi,t)$

$$\frac{\partial w(\psi,t)}{\partial t} = -\frac{\partial}{\partial \psi}\left[2\sqrt{I}w(\psi,t)\right] + K\nu(t)\left[\frac{I+V^2}{I+(V+g)^2}w(\psi^+,t) - w(\psi,t)\right] , \tag{S9}$$

where $\psi^+$ is the shifted phase:

$$V+g = \sqrt{I}\tan\frac{\psi^+}{2} , \qquad \tan\frac{\psi^+}{2} = \alpha + \tan\frac{\psi}{2} , \qquad \alpha \equiv \frac{g}{\sqrt{I}} = \frac{g_0}{\sqrt{i_0}K^{3/4}} .$$

By explicitating all the terms in the right-end side of Eq. (S9) in terms of the variable $\psi$

$$\frac{I+V^2}{I+(V+g)^2} = \frac{1+\tan^2\frac{\psi}{2}}{1+\left(\alpha+\tan\frac{\psi}{2}\right)^2} = \frac{1}{1+\frac{\alpha^2}{2}+\alpha\sin\psi+\frac{\alpha^2}{2}\cos\psi} ,$$

we can finally rewrite Eq. (S9) as

$$\frac{\partial w(\psi,t)}{\partial t} = -\frac{\partial}{\partial \psi}\left[2\sqrt{I}w(\psi,t)\right] + K\nu(t)\left[\frac{w(\psi_+,t)}{1+\frac{\alpha^2}{2}+\alpha\sin\psi+\frac{\alpha^2}{2}\cos\psi} - w(\psi,t)\right] . \tag{S10}$$

In Fourier space the PDF $w(\psi,t)$ can be expressed as

$$w(\psi,t) = \frac{1}{2\pi}\sum_{n=-\infty}^{+\infty} z_n e^{-in\psi},$$

with $z_0 = 1$ and $z_{-n} = z_n^*$, where the complex coefficients $z_n$ are the so-called Kuramoto–Daido order parameters $z_n$ [6, 7]. In terms of these coefficients the continuity equation (S10) becomes

$$\dot{z}_n = i2n\sqrt{I}z_n + K\nu(t)\left[\sum_{m=-\infty}^{+\infty} I_{nm}z_m - z_n\right] , \tag{S11}$$

where

$$I_{nm} \equiv \frac{1}{2\pi}\int_0^{2\pi} \frac{e^{in\psi}\left(e^{-i\psi^+}\right)^m \mathrm{d}\psi}{1+\frac{\alpha^2}{2}+\alpha\sin\psi+\frac{\alpha^2}{2}\cos\psi} . \tag{S12}$$

In order to calculate the integrals $I_{nm}$, we need to express explicitely the term $e^{-i\psi^+} = \cos\psi^+ - i\sin\psi^+$ entering in Eq. (S12). In particular, by noticing that

$$\cos\psi^+ = \frac{1-\tan^2\frac{\psi^+}{2}}{1+\tan^2\frac{\psi^+}{2}} , \qquad \sin\psi^+ = \frac{2\tan\frac{\psi^+}{2}}{1+\tan^2\frac{\psi^+}{2}} ,$$

and therefore that

$$e^{-i\psi^+} = \cos\psi^+ - i\sin\psi^+ = \frac{1-i\tan\frac{\psi^+}{2}}{1+i\tan\frac{\psi^+}{2}} = -\frac{\tan\frac{\psi}{2}+\alpha+i}{\tan\frac{\psi}{2}+\alpha-i} .$$

Moreover by expressing $\zeta \equiv e^{i\psi}$ as follows

$$\zeta \equiv e^{i\psi} = -\frac{\tan\frac{\psi}{2} - i}{\tan\frac{\psi}{2} + i} , \qquad \text{and hence} \quad \tan\frac{\psi}{2} = i\frac{1-\zeta}{1+\zeta} ,$$

one finally finds

$$e^{-i\psi_+} = -\frac{\alpha + 2i + \alpha\zeta}{\alpha + (\alpha - 2i)\zeta} . \tag{S13}$$

By employing the above expression the integral (S12) can be rewritten as

$$\begin{aligned} I_{nm} &= \int_0^{2\pi} \frac{(2\pi)^{-1} e^{in\psi} \left[-\frac{\alpha+2i+\alpha e^{i\psi}}{\alpha+(\alpha-2i)e^{i\psi}}\right]^m \mathrm{d}\psi}{1 + \frac{\alpha^2}{2} - \frac{i\alpha}{2}(e^{i\psi} - e^{-i\psi}) + \frac{\alpha^2}{4}(e^{i\psi} + e^{-i\psi})} = \oint_{|\zeta|=1} \frac{(i2\pi)^{-1}\zeta^{n-1}\left[-\frac{\alpha+2i+\alpha\zeta}{\alpha+(\alpha-2i)\zeta}\right]^m \mathrm{d}\zeta}{1 + \frac{\alpha^2}{2} - \frac{i\alpha}{2}(\zeta - \frac{1}{\zeta}) + \frac{\alpha^2}{4}(\zeta + \frac{1}{\zeta})} \\ &= \frac{1}{2\pi i} \oint_{|\zeta|=1} \frac{4(-\alpha)^m \zeta^n (\zeta + \frac{\alpha+2i}{\alpha})^{m-1}}{\alpha(\alpha-2i)^{m+1}(\zeta + \frac{\alpha}{\alpha-2i})^{m+1}} \mathrm{d}\zeta . \end{aligned} \tag{S14}$$

To evaluate the complex integral in (S14) we should identify the poles of the integrand. We will limit to consider $n \geq 1$ since $z_0 = 1$, and $z_{-n} = z_n^*$, for $m \leq 0$ the integrand has the following pole

$$\zeta_2 = -\frac{\alpha + 2i}{\alpha} ,$$

which is always beyond the unit circle since $|\frac{\alpha+2i}{\alpha}|^2 = 1 + 4/\alpha^2 > 1$ and, therefore, does not contribute to the integral $I_{nm}$. However, for $m \geq 0$ the integrand displays another pole located at

$$\zeta_1 = -\frac{\alpha}{\alpha - 2i} .$$

This pole is always within the integration contour $|\zeta| = 1$ since $|\frac{\alpha}{\alpha-2i}|^2 = \frac{\alpha^2}{\alpha^2+4} < 1$ and therefore it contributes to the integral (S14).

Let us first calculate the following integral

$$\begin{aligned} \frac{1}{2\pi i} \oint_{|\zeta|=1} \frac{\zeta^n (\zeta + \frac{\alpha+2i}{\alpha})^{m-1}}{(\zeta + \frac{\alpha}{\alpha-2i})^{m+1}} \mathrm{d}\zeta &= \begin{cases} \frac{1}{m!} \frac{\mathrm{d}^m}{\mathrm{d}\zeta^m} \left( \zeta^n \left(\zeta - \zeta_2\right)^{m-1} \right) \Big|_{\zeta=\zeta_1} , & m \geq 0 ; \\ 0 , & m \leq -1 \end{cases} \\ &= \begin{cases} \sum\limits_{j=0}^{\min(n,m)-1} \binom{m-1}{j} \binom{n+m-1-j}{m} \zeta_1^{n-1-j} \left(-\zeta_2\right)^j , & m \geq 1 ; \\ \frac{\zeta_1^n}{\zeta_1 - \zeta_2} , & m = 0 ; \\ 0 , & m \leq -1 \end{cases} \end{aligned}$$

where $\min(n, m)$ returns the minimal of two values, the binomial coefficients are defined as $\binom{n}{m} = \frac{n!}{m!(n-m)!}$. In particular, for obtaining the result reported in the latter line we considered separately the cases for $1 \leq n \leq m$ and for $n > m$. Hence,

$$I_{nm}(\alpha) = \begin{cases} \sum\limits_{j=0}^{\min(n,m)-1} \frac{(\zeta_1-\zeta_2)(n+m-1-j)! \cdot \zeta_1^{m+n-1-j} (-\zeta_2)^j}{m \cdot j! \cdot (m-1-j)! \cdot (n-1-j)!} , & m \geq 1 ; \\ \zeta_1^n , & m = 0 ; \\ 0 , & m \leq -1 . \end{cases} \tag{S15}$$

After the substitution of the values of $\zeta_1$ and $\zeta_2$ in (S15), the coefficients $I_{nm}(\alpha)$ take the form reported in Eq.(6) in the Letter.

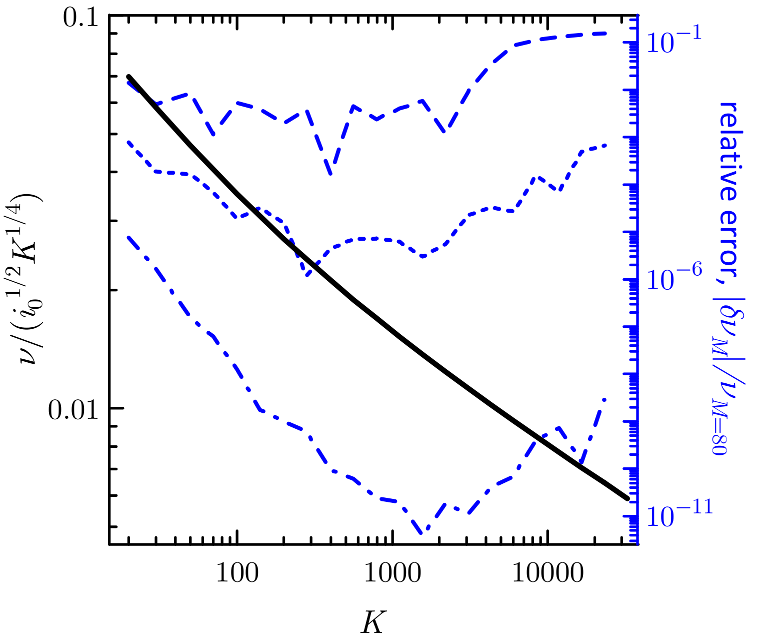


FIG. S2. The relative error in the estimation of the firing rate $\nu$ for a stationary situation computed with (S11)–(S16) plotted versus $K$ for $M = 10$ (dashed), 20 (dotted), 40 (dash-dotted). The reference firing rate was computed with $M = 80$ and it is shown in rescaled units as a bold solid line. Parameters: $(i_0, g_0) = (0.006, 1)$.

In particular, for $n \geq 1$, one has $I_{nm} = 0$ for $m < 0$; therefore, the matrix $I_{nm}$ has non zero elements only for $n = 1, 2, 3, ...$ and $m = 0, 1, 2, 3, ...$ :

$$\left(\frac{I_{nm}}{\zeta_1 - \zeta_2}\right) = \begin{pmatrix} \frac{\zeta_1}{\zeta_1-\zeta_2} & \zeta_1 & \zeta_1^2 & \zeta_1^3 & \dots \\ \frac{\zeta_1^2}{\zeta_1-\zeta_2} & 2\zeta_1^2 & 3\zeta_1^3 - \zeta_1^2\zeta_2 & 4\zeta_1^4 - 2\zeta_1^3\zeta_2 & \dots \\ \frac{\zeta_1^3}{\zeta_1-\zeta_2} & 3\zeta_1^3 & 6\zeta_1^4 - 3\zeta_1^3\zeta_2 & 10\zeta_1^5 - 8\zeta_1^4\zeta_2 + \zeta_1^3\zeta_2^2 & \dots \\ \frac{\zeta_1^4}{\zeta_1-\zeta_2} & 4\zeta_1^4 & 10\zeta_1^5 - 6\zeta_1^4\zeta_2 & 20\zeta_1^6 - 20\zeta_1^5\zeta_2 + 4\zeta_1^4\zeta_2^2 & \dots \\ & \dots & & & \end{pmatrix} .$$

In the continuity equation Eq. (S9) appears the population firing rate $\nu(t)$ that should be evaluated self-consistently during the time evolution. The istantaneous firing rate is fixed by the the flux at the firing threshold, therefore

$$\nu(t) = \lim_{V\to\infty} V^2 P(V,t) = 2\sqrt{I}\, w(\pi,t) = \frac{\sqrt{I}}{\pi}\mathrm{Re}\big(1 - 2z_1 + 2z_2 - 2z_3 + 2z_4 - \dots\big) \ . \qquad \text{(S16)}$$

corresponding to eq. (7) in the Letter.

*Accuracy of the numerical simulations of the system (S11,S16)*

The dynamical system (S11,S16) has been numerically integrated by truncating the the series $z_n$ at $n = M$. Usually we fix $M = 100$ and, where necessary, $M$ was elevated in order to keep the relative simulation error below $10^{-12}$ (see Fig. S2). In some cases the length of the digital mantissa of the variables was increased significantly beyond 16 digits in order to maintain this accuracy.

##### B. Linear stability and bifurcation analysis of the macroscopic evolution

In order to analyzie the stability of the asynchronous dynamics and the emergence of collective oscillations in the macroscopic evolution, we rewrite the system (S11,S16) as follows :

$$\begin{aligned}\frac{\mathrm{d}z_n}{\mathrm{d}t} &= F_n(z_1, z_2, z_3, \dots) \\ &= \sqrt{i_0}K^{\frac{1}{4}}\left(i2nz_n + K\widetilde{\nu}\Big[I_{n0}(\alpha) + \sum_{m=1}^{+\infty} I_{nm}(\alpha)\, z_m - z_n\Big]\right), \end{aligned} \tag{S17}$$

with

$$\widetilde{\nu} = \frac{1}{\pi}\mathrm{Re}(1 - 2z_1 + 2z_2 - 2z_3 + 2z_4 - \dots)\,. \tag{S18}$$

As a first aspect we notice that the dynamical evolution of the system (S17)–(S18) is controlled by two dimensionless parameters, namely

$$K \quad \text{and} \quad \alpha = \frac{g_0}{\sqrt{i_0}K^{3/4}}\,,$$

while the time can be rescaled by the factor $\sqrt{I} = \sqrt{i_0}K^{1/4}$. Therefore a bifurcation diagram in the plane $(K, i_0/g_0^2)$ is sufficient to capture all possible dynamical regimes observable for the system (S17)–(S18).

In order to analyze the stability of the stationary regimes, we should linearize the system (S17)–(S18) around a fixed point solution $\{z_n^{(0)}\}$. In particular, since $F_n$ depend on $\widetilde{\nu}$, which is non analytic function of $z_m$, the linearization of (S17) should be performed by considering as independent variables the real and complex part of the Kuramoto–Daido order parameters : namely, $z_n^{\mathrm{re}} = \mathrm{Re}(z_n)$ and $z_n^{\mathrm{im}} = \mathrm{Im}(z_n)$ (here and hereafter, superscripts "re" and "im" denote the real and imaginary parts).

Therefore the linearization of (S17) can be written as

$$\delta\dot{x}_{2n-1} = \frac{\partial F_n^{\mathrm{re}}}{\partial z_m^{\mathrm{re}}}\delta x_{2n-1} + \frac{\partial F_n^{\mathrm{re}}}{\partial z_m^{\mathrm{im}}}\delta x_{2n} \tag{S19}$$

$$\delta\dot{x}_{2n} = \frac{\partial F_n^{\mathrm{im}}}{\partial z_m^{\mathrm{re}}}\delta x_{2n-1} + \frac{\partial F_n^{\mathrm{im}}}{\partial z_m^{\mathrm{im}}}\delta x_{2n} \tag{S20}$$

where $\delta x_{2n-1}$ and $\delta x_{2n}$ are the infinitesimal perturbations of the real and imaginary parts of $z_n$, respectively. Moreover, the other terms entering in (S19) (S20) can be explicitly written as

$$\frac{\partial F_n^{\mathrm{re}}}{\partial z_m^{\mathrm{re}}} = \sqrt{i_0}K^{\frac{1}{4}}\left(K\widetilde{\nu}\big(I_{nm}^{\mathrm{re}} - \delta_{nm}\big) + K\frac{2(-1)^m}{\pi}\Big[I_{n0} + \sum_{m'=1}^{+\infty} I_{nm'}z_{m'}^{(0)} - z_n^{(0)}\Big]^{\mathrm{re}}\right), \tag{S21a}$$

$$\frac{\partial F_n^{\mathrm{im}}}{\partial z_m^{\mathrm{re}}} = \sqrt{i_0}K^{\frac{1}{4}}\left(2n\delta_{nm} + K\widetilde{\nu}I_{nm}^{\mathrm{im}} + K\frac{2(-1)^m}{\pi}\Big[I_{n0} + \sum_{m'=1}^{+\infty} I_{nm'}z_{m'}^{(0)} - z_n^{(0)}\Big]^{\mathrm{im}}\right), \tag{S21b}$$

$$\frac{\partial F_n^{\mathrm{re}}}{\partial z_m^{\mathrm{im}}} = \sqrt{i_0}K^{\frac{1}{4}}\left(-2n\delta_{nm} - K\widetilde{\nu}I_{nm}^{\mathrm{im}}\right)\,, \tag{S21c}$$

$$\frac{\partial F_n^{\mathrm{im}}}{\partial z_m^{\mathrm{im}}} = \sqrt{i_0}K^{\frac{5}{4}}\widetilde{\nu}\big(I_{nm}^{\mathrm{re}} - \delta_{nm}\big)\,. \tag{S21d}$$

The linear stability analysis of time-independent solutions $\mathbf{z}^{(0)} = \{z_n^{(0)}\}$ of system (S17)–(S18) can be performed by solving the eigenvalue problems associated to Eqs. (S19) (S20). In particular, by truncating the system (S17)–(S18) to the $M$-th Kuramoto-Daido order parameter the eigenvalue problem can be solved by diagonalizing a $(2M\times 2M)$ matrix with real-valued elements given by (S21) to find the associated complex eigenvalue spectrum $\{\lambda_k\} \quad k = 1, \dots, 2M$.

*Amplitude equations for a Hopf bifurcation*

As for the linear stability, also for the bifurcation analysis, one must handle $z_n^{\mathrm{re}}$ and $z_n^{\mathrm{im}}$ as independent real-valued variables. In particular, it is important to stress that the dynamical system (S17,S18) is quadratic with respect to the variables $z_n^{\rm re}$ and $z_n^{\rm im}$. The absence of cubic terms makes the calculation of the coefficients of the amplitude equations quite simple as we will see in the following.

Let us now analyse the Hopf instability of the stationary solution $\mathbf{z}^{(0)} = \{z_1^{(0)}, z_2^{(0)}, z_3^{(0)}, \dots\}$ in proximity of the critical value $K_{cr}$, where the in-degree $K$ is the control parameter and $i_0$ and $g_0$ are maintained constant. In order to perform this analysis we have considered *finite* perturbations $\mathbf{x} = \{x_1, x_2, x_3, x_4, \dots\}$ of the stationary solution, where $x_m$ are real-valued variables defined as follows: $z_n^{\rm re} \equiv (z_n^{(0)})^{\rm re} + x_{2n-1}$, $z_n^{\rm im} \equiv (z_n^{(0)})^{\rm im} + x_{2n}$. To obtain an *exact* evolution equation for the perturbations, we have substituted $z_n = z_n^{(0)} + x_{2n-1} + ix_{2n}$ into the system (S17,S18). Furthermore, by remembering that $\mathbf{z}^{(0)}$ solves the right-hand side of the system (S17,S18), and rearranging the terms, one can arrive to the following system of differential equations ruling the dynamics of $\mathbf{x}$:

$$\frac{{\rm d}x_n}{{\rm d}t} = F_{nm}x_m + \widetilde{F}_{nml}x_m x_l + (K - K_{\rm cr})\left(\frac{\partial F_{nm}}{\partial K}\right)_{i_0,g_0} x_m \,, \tag{S22}$$

where we imply the Einstein summation rule over repeated indices and odd- and even-order $x_n$ are the perturbations of the real and imaginary parts of $z_m$. Moreover, the coefficients $F_{nm}$ are given by (S21),

$$F_{2n-1\;2m-1} = \left.\frac{\partial F_n^{\rm re}}{\partial z_m^{\rm re}}\right|_{\mathbf{z}^{(0)}}, \qquad F_{2n\;2m-1} = \left.\frac{\partial F_n^{\rm im}}{\partial z_m^{\rm re}}\right|_{\mathbf{z}^{(0)}},$$
$$F_{2n-1\;2m} = \left.\frac{\partial F_n^{\rm re}}{\partial z_m^{\rm im}}\right|_{\mathbf{z}^{(0)}}, \qquad F_{2n\;2m} = \left.\frac{\partial F_n^{\rm im}}{\partial z_m^{\rm im}}\right|_{\mathbf{z}^{(0)}};$$

while the quadratic form coefficients $\widetilde{F}_{nml}$ come from the term

$$K\left(\widetilde{\nu} - \pi^{-1}\right)\sum_{m=1}^{+\infty} I_{nm} z_m^{(0)} = \sum_{m,m'} \left.\frac{2K(-1)^{m'}}{\pi} I_{nm} z_m z_{m'}^{\rm re}\right|_{\mathbf{z}^{(0)}},$$

which yields

$$\widetilde{F}_{2n-1\;ml}x_m x_l = \sum_{m,m'} \left.\frac{2\sqrt{i_0}K^{\frac{5}{4}}(-1)^{m'}}{\pi}\left[I_{nm}^{\rm re} z_m^{\rm re} - I_{nm}^{\rm im} z_m^{\rm im}\right] z_{m'}^{\rm re}\right|_{\mathbf{z}^{(0)}},$$
$$\widetilde{F}_{2n\;ml}x_m x_l = \sum_{m,m'} \left.\frac{2\sqrt{i_0}K^{\frac{5}{4}}(-1)^{m'}}{\pi}\left[I_{nm}^{\rm re} z_m^{\rm im} + I_{nm}^{\rm im} z_m^{\rm re}\right] z_{m'}^{\rm re}\right|_{\mathbf{z}^{(0)}};$$

and $\left(\frac{\partial \dots}{\partial K}\right)_{i_0,g_0}$ indicates the derivative with respect to $K$ for fixed $i_0$, $g_0$.

At the threshold of a Hopf instability, the most unstable modes are associated to two purely imaginary and complex conjugates eigenvalues. Therefore the linear stability matrix $(F_{nm})$ possesses a pair of eigenvalues $\pm i\Omega \equiv \pm i2\pi f^H$ with eigenvector $\mathbf{Y}$ and $\mathbf{Y}^*$, namely:

$$i\Omega Y_n = F_{nm}Y_m\,, \qquad -i\Omega Y_n^* = F_{nm}Y_m^*\,, \tag{S23}$$

and the Hermitian conjugate problem possesses eigenvalues $\mp i\Omega$ with eigenvectors $\mathbf{Y}^+$ and $(\mathbf{Y}^+)^*$ :

$$-i\Omega Y_n^+ = F_{mn}Y_m^+\,, \qquad i\Omega (Y_n^+)^* = F_{mn}(Y_m^+)^*\,. \tag{S24}$$

To find the amplitude equation we employ the standard multiple scale method [6] with formal small parameter $\epsilon$; $\frac{\rm d}{{\rm d}t} = \frac{\partial}{\partial t_0} + \epsilon\frac{\partial}{\partial t_1} + \epsilon^2\frac{\partial}{\partial t_2} + \dots$, $x_n = \epsilon x_n^{(1)} + \epsilon^2 x_n^{(2)} + \epsilon^3 x_n^{(3)} + \dots$, $K = K_{\rm cr} + \epsilon^2 K_2 + \dots$, where $t_0$ is the "normal" time and $t_1$, $t_2$ are the "slow" times. By substituting these expansions in Eq. (S22) we obtain at the first order in $\epsilon$ the following expression

$$x_n^{(1)} = A(t_1, t_2, \dots)Y_n e^{i\Omega t_0} + c.c.\,.$$

And at the order $\epsilon^2$:

$$\frac{\partial x_n^{(2)}}{\partial t_0} + \frac{\partial A}{\partial t_1}Y_n e^{i\Omega t_0} + c.c. = F_{nm}x_m^{(2)} + \widetilde{F}_{nml}\left[A^2 Y_m Y_l e^{i2\Omega t_0} + c.c. + |A|^2(Y_m Y_l^* + Y_m^* Y_l)\right].$$

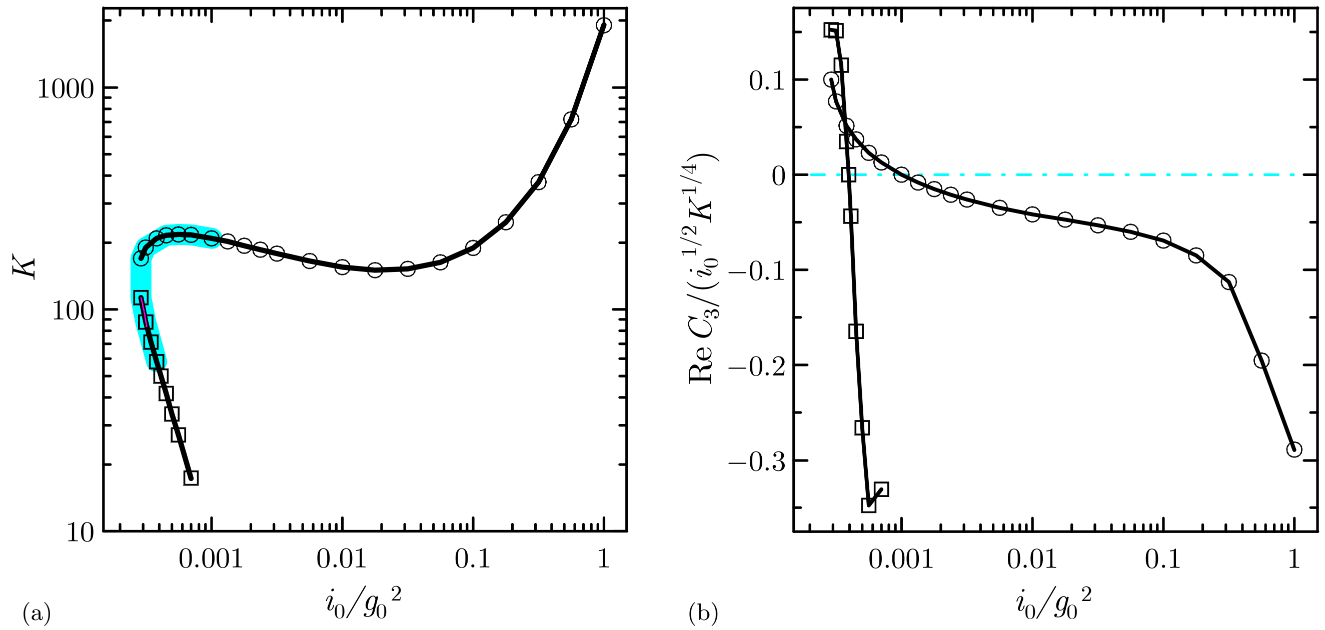


FIG. S3. Phase diagram of the macroscopic behavior of a QIF population (S17,S18). (a) Hopf bifurcation lines in the plane $(i_0/g_0^2, K)$. The cyan shading identifies the region where the bifurcations are subcritical. (b) The coefficient $\mathrm{Re}\, C_3$ in rescaled units at the critical points $K = K_{cr}$ versus $i_0/g_0^2$. In both panels circles and squares denote the two possibly coexisting bifurcation branches for fixed values of $i_0/g_0^2$.

One finds $\partial A/\partial t_1 = 0$ and

$$x_n^{(2)} = A^2 \chi_n e^{i2\Omega t_0} + c.c. + |A|^2 \phi_n \,,$$

where $\phi_n$ is the solution of the linear equation system

$$F_{nm}\phi_m = -\widetilde{F}_{nml}(Y_m Y_l^* + Y_m^* Y_l) \,,$$

and $\chi_n$ is the solution of

$$(F_{nm} - i2\Omega\delta_{nm})\chi_m = -\widetilde{F}_{nml} Y_m Y_l \,.$$

At the order $\epsilon^3$, we find the solvability condition,

$$\frac{\partial A}{\partial t_2}(Y_{n'}^+)^* Y_{n'} = K_2 A (Y_n^+)^* \left(\frac{\partial F_{nm}}{\partial K}\right)_{i_0,g_0} Y_m + A|A|^2 (Y_n^+)^* \widetilde{F}_{nml} \left(Y_m^* \chi_l + \chi_m Y_l^* + Y_m \phi_l + \phi_m Y_l\right) \,.$$

The latter is the so-called amplitude equation and it can be recast in the standard form :

$$\frac{\partial A}{\partial t} = (K - K_{\mathrm{cr}}) \frac{\mathrm{d}\lambda}{\mathrm{d}K} A + C_3 A|A|^2 \,, \tag{S25}$$

where $A$ is the complex amplitude of oscillations, $x_n = A Y_n e^{i\Omega t} + c.c. + \mathcal{O}(A^2)$; $\lambda$ is the exponential growth rate of the leading mode. The expression of the coefficient $C_3$ being the following

$$C_3 = \frac{(Y_n^+)^* \widetilde{F}_{nml} \left(Y_m^* \chi_l + \chi_m Y_l^* + Y_m \phi_l + \phi_m Y_l\right)}{(Y_{n'}^+)^* Y_{n'}} \,. \tag{S26}$$

The sign of the real part of $C_3$ identifies the Hopf bifurcation as sub-critical (super-critical) for Re $C_3 > 0$ (Re $C_3 < 0$).

The Hopf bifurcation lines in the plane $(i_0/g_0^2, K)$ obtained with this approach are reported in Fig. S3 (a), the bifurcation are always super-critical aparte in the cyan shaded region, where they are sub-critical. The sub- or super-critical nature of the Hopf bifurcation is decided on the basis of the sign of Re $C_3$ displayed in Fig. S3 (b) as a function of $i_0/g_0^2$ at the critical points $K = K_{cr}$.

[1] A. Tonnelier, H. Belmabrouk, and D. Martinez, Neural Computation **19**, 3226 (2007).
[2] A. Torcini and P. Politi, The European Physical Journal B-Condensed Matter and Complex Systems **25**, 519 (2002).
[3] M. di Volo and A. Torcini, Phys. Rev. Lett. **121**, 128301 (2018).
[4] G. B. Ermentrout and N. Kopell, SIAM Journal on Applied Mathematics **46**, 233 (1986).
[5] B. Ermentrout, Scholarpedia **3**, 1398 (2008), revision #122134.
[6] Y. Kuramoto, *Chemical oscillations, waves, and turbulence*, Vol. 19 (Springer Science & Business Media, 2012).
[7] H. Daido, Progress of theoretical physics **88**, 1213 (1992).
